# Expression profile of CASSIOPEIA patients refines prognostic value of MRD negativity in multiple myeloma

**DOI:** 10.64898/2026.04.07.716874

**Authors:** Florence Magrangeas, Catherine Guérin-Charbonnel, Victor Bessonneau-Gaborit, Marie Denoulet, Nils Giordano, Aurore Perrot, Cyrille Touzeau, Mark van Duin, Magali Devic, Elise Douillard, Eric Letouzé, Peter Sonneveld, Jill Corre, Stéphane Minvielle, Philippe Moreau

## Abstract

Long-term follow-up of the CASSIOPEIA trial (NCT02541383) demonstrated superior progression-free survival (PFS) with daratumumab, both in combination with bortezomib, thalidomide, and dexamethasone during induction and consolidation, and during maintenance therapy, in transplant- eligible patients newly diagnosed with multiple myeloma (MM). However, outcomes among CASSIOPEIA patients remain heterogeneous across treatment groups. Measurable residual disease (MRD) is a strong indicator of the depth and duration of therapeutic response and is independently associated with both PFS and overall survival (OS), but it does not fully capture the biological diversity of MM. We performed a risk prediction analysis based on transcriptomic subgroups in CASSIOPEIA patients. A subset of 628 patients was characterized using RNA sequencing and consensus clustering identified five transcriptomic subtypes of MM. Long-term follow-up allowed the definition of three transcriptomic risk categories, with estimated 72-month PFS rates of 70%, 51%, and 27% for low, intermediate, and high-risk groups, respectively, among patients who received daratumumab in at least one treatment phase. In these patients, MRD negativity rates after consolidation and six months later were significantly higher in the low and high-risk groups compared with the intermediate-risk group. In the high-risk group, MRD status was not associated with PFS or OS. This suggests that, although daratumumab administered during both the induction/consolidation and maintenance phases improves the clinical outcomes of patients with activation of NSD2 or overexpressing members of the MAF family, highly aggressive minor clones may rapidly expand. These findings emphasize the need for novel therapeutic strategies in this high-risk population.

## Introduction

Multiple myeloma (MM) remains a challenging hematologic malignancy, although significant therapeutic advances have markedly improved patient outcomes. Among these, anti-CD38 monoclonal antibodies such as daratumumab^1^ have revolutionized frontline therapy by enhancing response rates and prolonging progression-free survival (PFS).^2–9^ The phase 3 randomized CASSIOPEIA trial (NCT0254138) evaluated the impact of adding daratumumab to bortezomib, thalidomide, and dexamethasone (VTd) both before and after autologous stem cell transplantation (part 1, first randomization), as well as the efficacy of daratumumab maintenance therapy for two years versus observation (part 2, second randomization) in patients with newly diagnosed multiple myeloma (NDMM).^6–8^ The trial demonstrated that the addition of daratumumab during induction and consolidation significantly improved both PFS and overall survival (OS), and that daratumumab maintenance further enhanced patient outcomes. However, daratumumab did not overcome the poor prognosis associated with high-risk cytogenetic abnormalities, namely del(17p) and t(4;14). The study also revealed that patients achieving measurable residual disease (MRD) negativity after induction, consolidation, and during maintenance experienced superior outcomes, with sustained MRD negativity correlating with prolonged PFS. These findings led to the approval of daratumumab-VTd (D-VTd) by both the EMA and FDA. Recently, the PERSEUS trial confirmed the benefit of adding subcutaneous daratumumab to bortezomib, lenalidomide and dexamethasone (VRd) in transplant-eligible patients, leading to a 58% reduction in the risk of progression or death.^9^ In the updated analysis with a follow-up exceeding 80 months, we investigated, using RNA sequencing (RNA-seq) data from CASSIOPEIA patients, the relationship between genomic subgroups, post-consolidation MRD status, and survival outcomes.

## Methods

### Biocollection and nucleic acid purification

Our population consists of 628 patients enrolled in the CASSIOPEIA trial (ClinicalTrials.gov identifier NCT02541383) for whom bone marrow samples collected at diagnosis provided enough CD138+ myeloma cells to perform transcriptomic analysis. RNA-seq data for 628 patients and DNA-seq data for 340 patients were generated from plasma cells sorted of bone marrow with automated magnetic sorting targeting CD138+ cells (autoMACS and anti-CD138 beads from Miltenyi Biotec, Bergisch Gladbach, Germany). Mean purity of CD138+ cells: 98.1% (60-100), RNA and DNA samples were isolated with AllPrep DNA/RNA/miRNA Universal Kit (Qiagen, Hilden, Germany) .

### RNA-seq library preparation

Input RNA quantity was based on qubit quantification with the Qubit RNA BR Assay kit (Invitrogen, Q10211) and RNA quality was checked using an Agilent bioanalyzer with the RNA 6000 Nano kit (Agilent, 5067-1511). Samples with an average RNA integrity number of 9 (range, 5.8-9.9) were processed with an average amount of 200 ng RNA input (range, 100-200 ng). RNA-seq libraries were constructed using the NEBNext Poly(A) mRNA magnetic isolation module (New England Biolabs, E7490L) and the NEBNext Ultra II Directional RNA Library Prep with Sample Purification Beads (New England Biolabs, E7765L) following the manufacturer’s recommendations. Final libraries were evaluated on the Agilent TapeStation using the high-sensitivity ScreenTape D1000 (Agilent, 5067-5584, 5067-5585) and quantified using the NEBNext Library Quant Kit for Illumina (New England Biolabs, E7630L). Libraries were sequenced on NovaSeq 6000 (Illumina, San Diego, CA) using paired-end 2*100 cycles sequencing.

### High genetic risk markers identification

t(4;14) (≥30% abnormal cells) or del(17p) (≥50% abnormal cells) chromosomal abnormalities were identified by fluorescence in situ hybridization during screening; del(1p32), 1q gain and *TP53* mutation were assessed by whole exome sequencing in 379 patients of the RNA-seq cohort. Biallelic del(1p32) was not considered separately as it was only identified in one patient

### MRD status

For all patients with MRD data available for a considered timepoint, MRD status was assessed using standardized EuroFlow-based multiparametric flow cytometry (MFC) at the 10^-5^ sensitivity threshold.

### Primary RNA-seq data analysis

FASTQ files were first trimmed to remove adapter sequences using Trimmomatic (v0.38). Trimmed reads were aligned to the human reference genome (hg19, release 75) using STAR (v2.5.3a) and reads with low mapping quality (MAPQ < 15) were filtered out using Samtools (v1.4.1). Raw counts were generated using featureCounts (v1.6.3) from the Subread suite, with annotation provided by the Ensembl GTF file (GRCh37, release 87).

### Secondary analysis procedures

#### Selection of variables

A filter was applied to retain only genes expressed at 1 count-per-million (cpm) for at least 1% of the population, and library sizes were recalculated. Count data were then normalized using the “trimmed-mean of the M-values” (TMM) method from the R package edgeR (edgeR_3.36.0). Dispersion values (negative binomial distribution) of the genes were then estimated using the “estimateDisp” function, and the ratio of the actual dispersion to the dispersion predicted by a local regression based on the mean of the log-expression was computed, with the addition of the constant 1 to the denominator to avoid a strong impact of small dispersions.

The non-parametric definition of outlier values was used to identify genes with higher-than-expected dispersion: “outlier genes”. Only outlier genes encoding proteins were subsequently considered. As a result, 932 outlier genes were identified. Then, based on log-expressions of these genes (log2(TMM+1)), the consensus clustering method was applied using the M3C package (M3C_1.16.0). *Differential analysis*

Genes with significant differential expression between one cluster and the others were identified with function exactTest of package edgeR. For each cluster, pathways corresponding to the most differential genes (FDR < 1e^-7^) were investigated using the enrichR package (enrichR_3.2), specifically the ‘MSigDB_Hallmark_2020’ module.

### Statistical analysis

Quantitative parameters were described by median, range, mean and standard deviation (sd) and compared with Welch’s test, along with Tukey HSD test in case of 3 or more groups compared, Wilcoxon test or Kruskal-Wallis test, as appropriate. Associations between genes were assessed by Spearman’s correlation coefficients. Qualitative parameters were described by number and percentage of each modality and compared with Chi-square test or Fisher exact test, as appropriate. Hierarchical clustering (patients or genes) was based on correlation distance and Ward’s method. Survival outcomes were estimated by Kaplan-Meier method and compared with logrank test. Hazard ratios, along with their 95% confidence intervals (CI), were assessed by univariable and multivariable Cox models. P-values lower than 5% were considered significant. P-values lower than 10% were considered as tendency.

All analyses were performed with R Software, version 4.1.3 (R Core Team (2014). R: A language and environment for statistical computing. R Foundation for Statistical Computing, Vienna, Austria. URL: http://www.R-project.org/).

## Results

### CASSIOPEIA RNA-seq cohort description

The CASSIOPEIA study enrolled 1,085 patients who were randomly assigned in induction and consolidation phase to the D-VTd group or to the VTd group, then 886 patients who completed consolidation and achieved at least a partial response were re-randomized to daratumumab maintenance or observation (**Figure 1A**). We conducted an ancillary analysis of the CASSIOPEIA study using an RNA-seq dataset generated from CD138⁺-purified bone marrow samples of 628 patients (**Supplemental Figure 1**). Among these, 288 patients in the D-VTd group and 301 in the VTd group completed consolidation therapy, after which 490 patients were re-randomized to receive either daratumumab maintenance (n = 239) or observation (n = 251). At the time of data cut-off, the median follow-up was 81.7 months (95% CI, 80.9–82.7) from the first randomization and 72.4 months (95% CI, 71.7–73.6) from the second randomization. Baseline characteristics of the transcriptomic (RNA-seq) cohort were comparable to those of the overall study population, except for differences in the distribution of ISS stages (**Supplemental Table 1**). Consistent with the findings in the full cohort, both PFS and OS from the first randomization, regardless of the second randomization, were significantly longer in the D-VTd group compared with the VTd group. Median PFS was 83.6 months (95% CI, 65.1– NR) in the D-VTd group versus 50.6 months (95% CI, 41.7–59.5) in the VTd group (HR, 0.6; 95% CI, 0.5– 0.7; *p* < 0.0001). For OS, the HR was 0.5 (95% CI, 0.3–0.7; *p* = 0.0002), with median OS not reached in either arm (**Supplemental Figure 2A–B**). Among the 490 patients who underwent the second randomization, 117 from the D-VTd arm received daratumumab maintenance, 129 were assigned to observation, 122 from the VTd arm received daratumumab maintenance, and 122 were assigned to observation. PFS was significantly longer in the D-VTd+daratumumab, D-VTd+observation, and VTd+daratumumab groups compared with the VTd+observation group (HRs 0.3 [95% CI, 0.2–0.5], 0.4 [95% CI, 0.3–0.6], and 0.4 [95% CI, 0.3–0.5], respectively; *p* < 0.0001; **Supplemental Figure 2C**). OS was also significantly longer in patients receiving D-VTd+daratumumab compared with VTd+observation (HR, 0.3; 95% CI, 0.2–0.7; *p* = 0.0024), and in those in the D-VTd+observation group versus VTd+observation (HR, 0.5; 95% CI, 0.3–0.9; *p* = 0.0257). Furthermore, among patients randomized to daratumumab maintenance, outcomes differed significantly according to prior exposure to daratumumab during induction and consolidation (HR, 0.4; 95% CI, 0.2–0.8; *p* = 0.0073; **Supplemental Figure 2D**). Since PFS and OS benefits associated with daratumumab in the quadruplet regimen were consistent between the RNA-seq cohort and the overall population, this genomic ancillary study provided a unique opportunity to explore the impact of daratumumab according to genomic subgroups.

**Figure 1.**
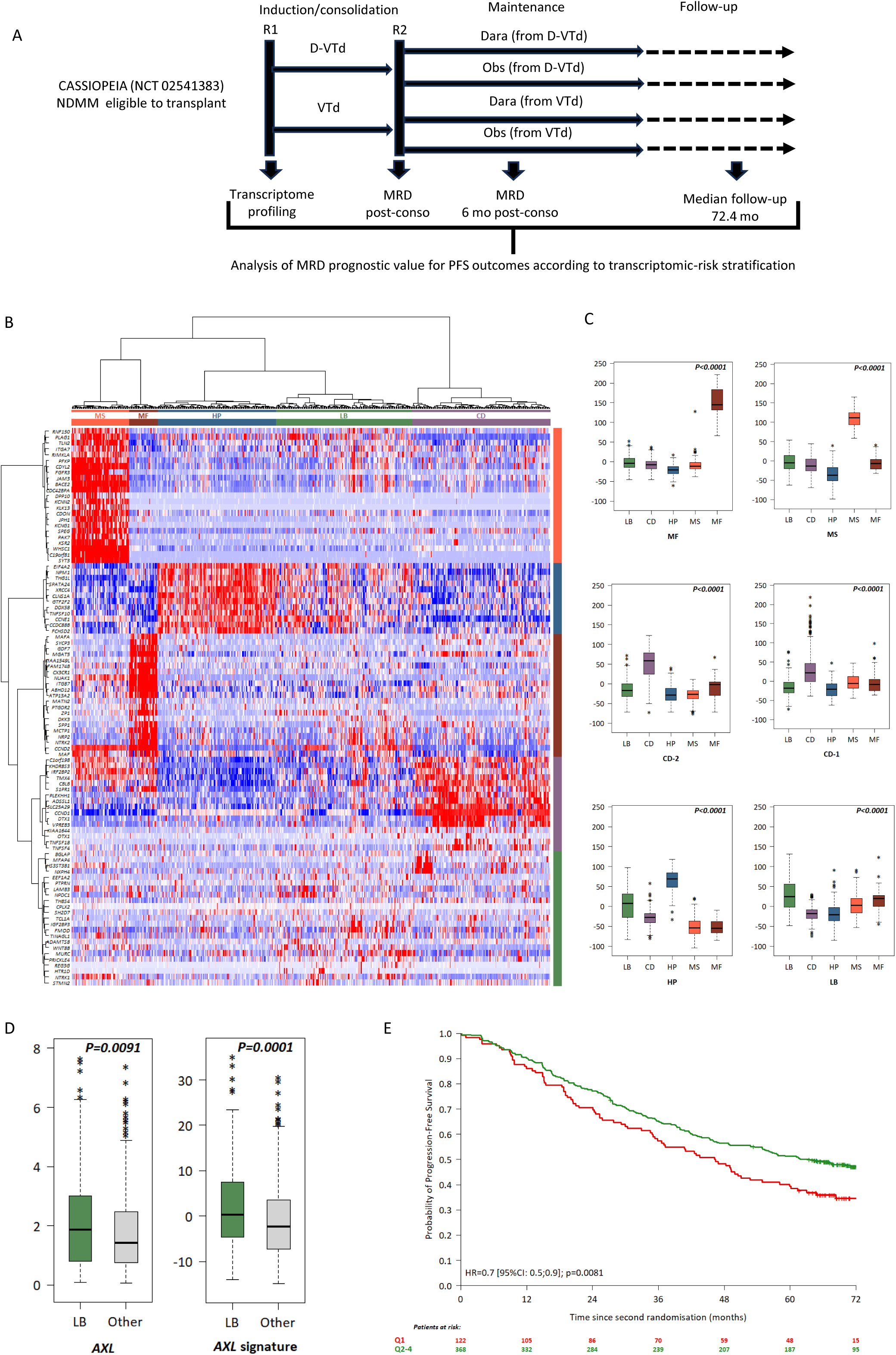
Gene expression subtypes in myeloma cells from CASSIOPEIA RNA-seq cohort. (A) Study design. (B) Unsupervised hierarchical clustering of the 932 outlier genes expression data of the CASSIOPEIA RNA-seq cohort. Top, the sample dendrogram identified five RNA subtypes, bottom heatmap showed expression of the 20 most differentially expressed genes between the RNA subtypes (95 unique genes). (C) Boxplots displaying distribution of the scores for the MS, MF, CD1, CD2, HP, LB and PR classification groups established by Zhan et al^10^ according to the 5 RNA subtypes. (D) Boxplots displaying *AXL* (left) and *AXL*-coregulated genes signature (right) for LB patients vs. patients in the 4 other subtypes. (E) Kaplan-Meier curves and Cox model results for PFS with *AXL*-coregulated genes signature dichotomized values (Q1 vs. Q2-4).

### Transcriptomic subtypes identification

To characterize the genomic subgroups of MM, we analyzed RNA-seq profiles from 628 patients enrolled in the CASSIOPEIA study. Unsupervised consensus clustering identified five distinct molecular clusters (**Figure 1B; Supplemental Figure 3**). Three clusters were defined by the overexpression of canonical target genes associated with immunoglobulin translocations. The *NSD2*-expressing cluster, characterized by t(4;14) and showing >97% concordance with FISH, corresponded to the previously described MS subtype^10^ and comprised 76 patients (12.1%). The *MAF/MAFA/MAFB*-expressing cluster, known as the MF subtype, included 37 patients (5.9%). The *CCND1*-expressing cluster, encompassing both CD-1 and CD-2 subtypes and referred to as CD, comprised 181 patients (28.8%). The fourth cluster, closely related to hyperdiploid (HP)-associated subtypes, included 155 patients (26.5%), while the fifth cluster, consisting of 179 patients (28.5%), corresponded to the so-called “low bone” (LB) subtype (**Figure 1C**). Notably, none of these five RNA-defined subtypes corresponded to the previously identified proliferation subtype (**Supplemental Figure 4**).^10^ Gene ontology (GO) pathway enrichment analyses comparing each RNA subtype against all others revealed numerous significantly enriched pathways. In the MF subtype, enriched pathways included IL-2/STAT5 signaling, Hedgehog signaling, mTORC1 signaling, and TNF-α signaling via NF-κB, suggesting a global transcriptional dysregulation driven by activation of the MAF transcription factor family. In contrast, the HP subtype exhibited enrichment for pathways related to stress adaptation, such as unfolded protein response, oxidative phosphorylation, and DNA repair likely reflecting the increased replicative burden associated with genomic gains. Additional enrichment of interferon response and MYC target pathways, known to be deregulated in MM with extensive copy-number alterations, was also observed (**Supplemental Figure 5**).^11^ Since no distinct proliferative RNA subtype was identified, we next investigated which subtypes displayed higher proliferative activity.^12–13^ Both MS and MF subtypes showed a significantly greater proportion of proliferating cells compared with the HP subtype (*P* = 0.0062 and *P* = 0.0117, respectively). Conversely, we examined which subtypes were enriched in non-proliferating, or dormant, myeloma cells.^14^ The *AXL* gene, a prototypical marker of dormancy, along with nine co-regulated genes (including *VCAM1*, *FCER1G*, *MPEG1*, and *CSF1R*) comprising the dormancy signature^15^ were significantly upregulated in the LB cluster compared with other subtypes (*P* = 0.0001; **Figure 1D; Supplemental Figure 6**). Furthermore, patients with low expression (Q1) of the *AXL* co-regulated dormancy gene signature exhibited shorter PFS (HR, 0.7 [95% CI, 0.5–0.9]; *P* = 0.0081; **Figure 1E**) as previously reported by Khoo *et al.*^15^

### PFS and OS according to transcriptomic subtypes

Given that CASSIOPEIA patients were classified into five distinct molecular entities, we hypothesized that each entity might confer different clinical outcomes. Among the 490 patients from the RNA-seq cohort who reached the maintenance phase, marked differences in outcomes were observed across molecular subtypes. Patients belonging to the MF and MS subtypes exhibited the poorest PFS compared with other groups, whereas HP and CD subtypes showed inferior PFS relative to the LB subtype (**Figure 2A**). Pairwise comparisons of outcomes among the subtypes enabled the identification of three prognostically distinct risk categories. The low-risk group, consisting of the LB subtype, had a median PFS not reached (NR; 95% CI, 80.5–NR). The intermediate-risk group, combining CD and HP subtypes, displayed a median PFS of 54.9 months (95% CI, 45.4–66.1), while the high-risk group, comprising the MS and MF subtypes, had a median PFS of 28.9 months (95% CI, 22.0–38.3) (**Figure 2B**). OS was significantly longer in the low- and intermediate-risk groups compared with the high-risk group (HR = 0.5 [95% CI, 0.3–0.8] and 0.3 [95% CI, 0.2–0.5], respectively; *P* < 0.0001; **Figure 2C**). As expected, patients who received daratumumab during induction and consolidation (n = 368) experienced superior PFS outcomes, although differences persisted among the three risk groups. Median PFS was not reached for low-risk patients, 74.2 months (95% CI, 55.7–NR) for intermediate-risk patients, and 33.0 months (95% CI, 27.6–43.9) for high-risk patients (**Figure 2D**). OS was also significantly longer in low- and intermediate-risk groups compared with the high-risk group (HR = 0.4 [95% CI, 0.2–0.7]; *P*=0.0010). Regarding the cytogenetic abnormalities included in the revised high-risk MM definition^16^ available in this study, aside from the t(4;14) translocation, which characterizes high-risk MM patients with MS, both 1q gain and del(1p32) were enriched in the high-risk group and, to a lesser extent, in the low-risk group. In contrast, del(17p) with a cutoff of > 20%, as well as *TP53* mutation, were evenly distributed across risk categories (**Supplemental Figure 7**). Notably, del(17p) was strongly associated with inferior OS (*p* = 0.0008) only within the intermediate-risk group.

**Figure 2.**
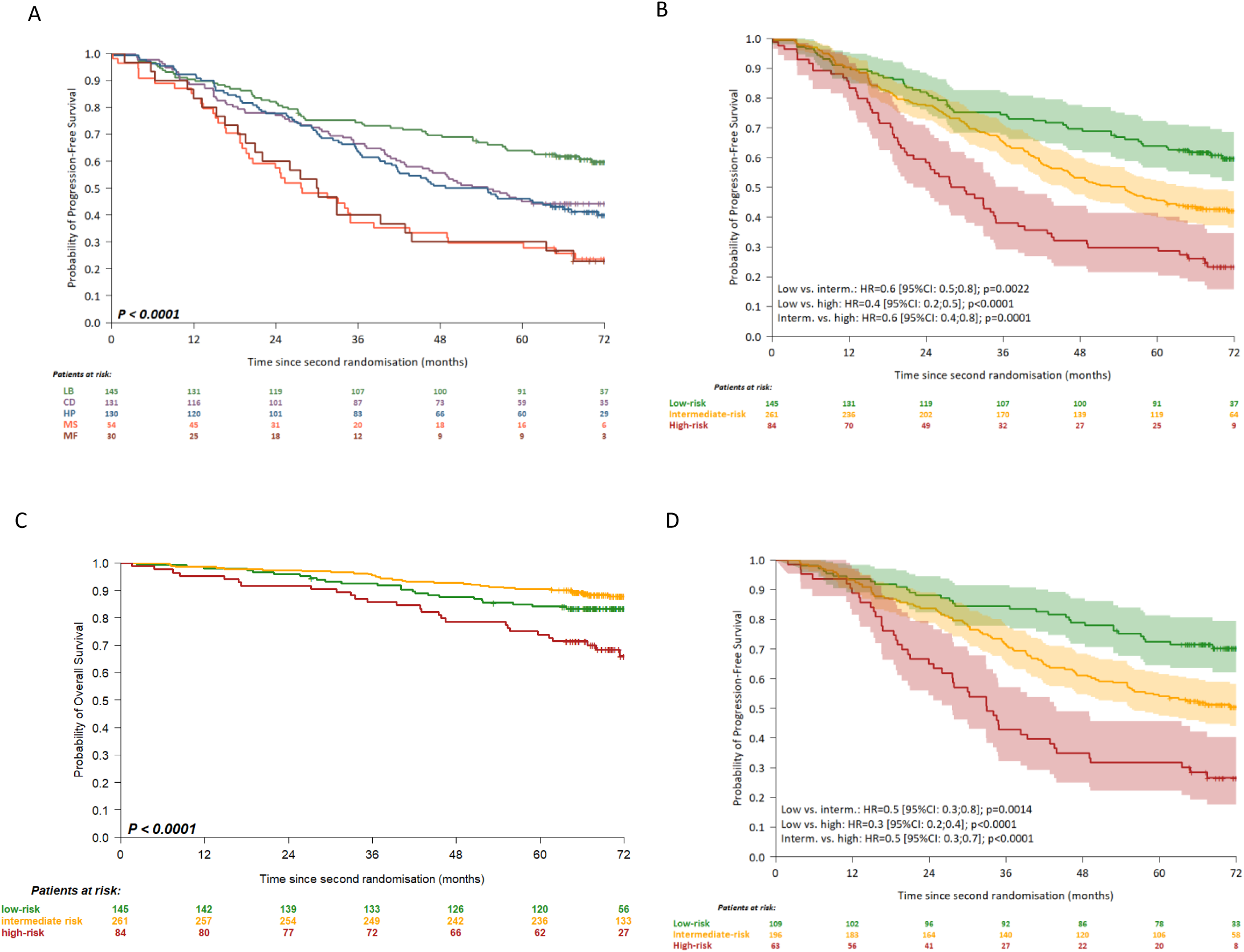
**Outcome association with transcriptomic subtypes**. (A) PFS according to the 5 RNA subtypes: LB, CD, HP, MS and MF. (B) Pairwise outcome comparison according the 3 risk groups: Low (LB), Intermediate (CD and HP), High (MS and MF). (C) Overall survival according to risk status: low-risk or intermediate-risk *vs.* high-risk. (D) PFS according to risk status in patients who have received daratumumab in at least one of the treatment phases.

### Clinical benefit of daratumumab according to transcriptomic risk status

We next evaluated the added benefit of daratumumab according to molecular risk group. Low- and intermediate-risk patients derived the greatest benefit from daratumumab, irrespective of the treatment phase during which the anti-CD38 antibody was administered (*P* < 0.0001; **Figure 3A–B**). In contrast, among high-risk patients, those who received daratumumab during both induction/consolidation and maintenance phases experienced a significantly improved PFS compared with patients who never received daratumumab (*P* = 0.0082) or those treated with daratumumab only during induction and consolidation (*P* = 0.0174) (**Figure 3C**). The latter group displayed a PFS comparable to that of patients who never received daratumumab (*P* = 0.61). These differences could not be attributed to variations in *CD38* expression across treatment arms (**Supplemental Figure 8**).

**Figure 3.**
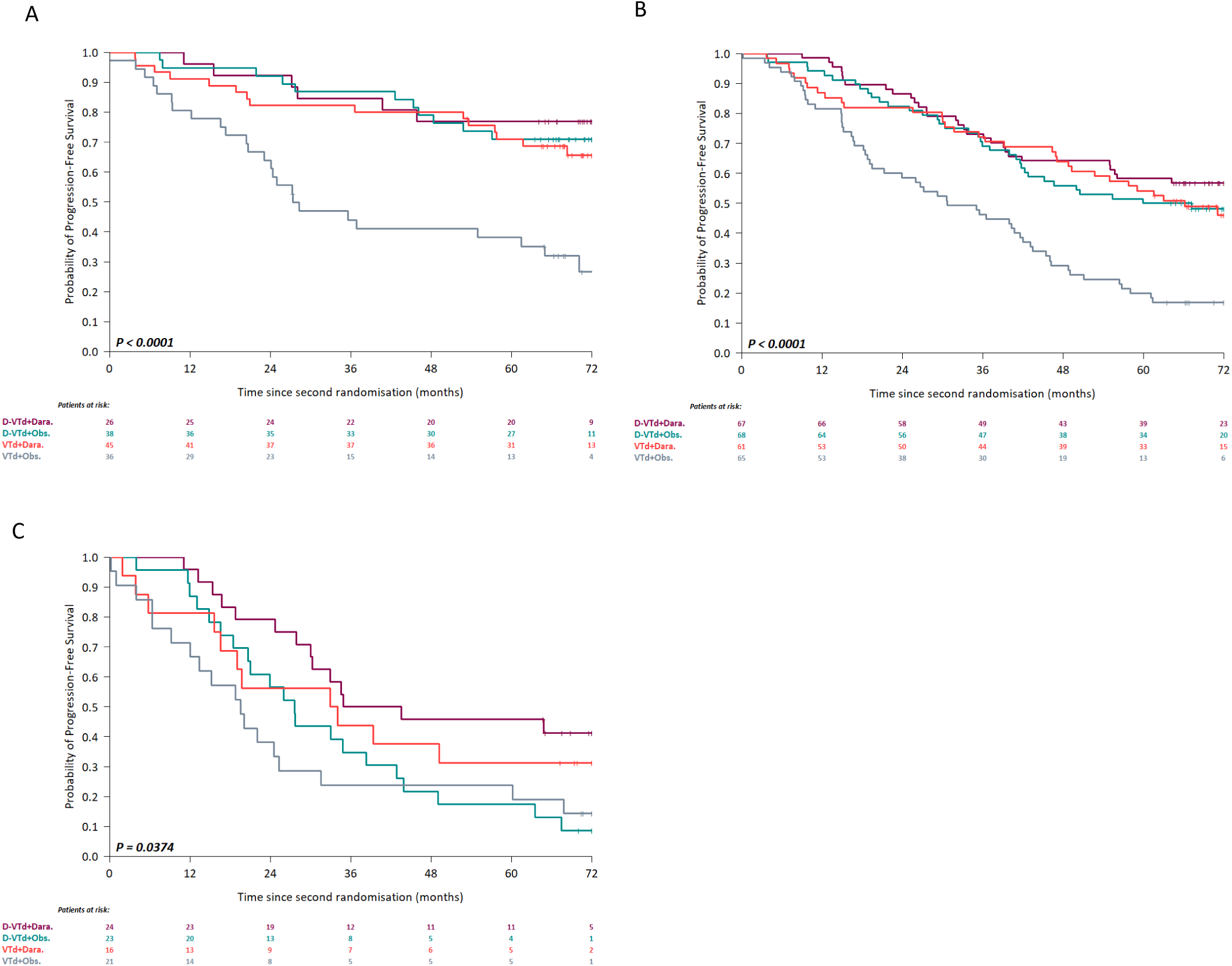
Daratumumab benefit according to RNA risk groups. (A) Low-risk patients, PFS estimates are shown for each treatment arm. Median PFS was reached only for VTd + Obs. (27.4 months, 95% CI, 23.1-70.1). Pairwise comparisons identified two significantly different subgroups D-VTd + Dara., D-VTd + Obs., VTd + Dara. and VTd + Obs. (*P*< 0.0001). (B) Intermediate-risk patients, PFS estimates are shown for each treatment arm. Median PFS was reached only for VTd + Obs. (30.7 months, 95% CI, 21.2-43.1). Pairwise comparisons identified two significantly different subgroups D-VTd + Dara., D-VTd + Obs., VTd + Dara. and VTd + Obs. (*P*< 0.0001). (C) High-risk patients, PFS estimates are shown for each treatment arm. Median PFS was reached for all treatment arms: VTd + Obs. (19.5 months, 95% CI, 12.0-60.2); VTd + Dara. (33.5 months, 95% CI, 16.6-NR); D-VTd +Obs. (27.6, months, 95% CI, 20.6-43.9) and D-VTd + Dara. (39.2 months, 95% CI, 30.2-NR). Pairwise comparisons identified two significantly different subgroups D-VTd + Dara. and VTd +Obs. (p=0.0082) or D-VTd +Obs. (P= 0.0174).

### Impact of *CD38* gene expression on PFS

The prognostic significance of baseline *CD38* expression levels in NDMM patients treated with daratumumab-containing quadruplet regimens remains unclear. To address this, we analyzed the relationship between *CD38* gene expression and clinical outcomes in CASSIOPEIA patients. Higher *CD38* expression was associated with prolonged PFS among patients who received daratumumab, independent of treatment phase, molecular risk group, or both (**Figure 4A; Supplemental Figure 9**). As expected, *CD38* expression had no impact on PFS in patients who did not receive daratumumab. Interestingly, the inclusion of daratumumab during both induction/consolidation and maintenance phases, or during maintenance alone, appeared to mitigate the prognostic effect of high *CD38* expression (**Figure 4B**). We further examined the interaction between molecular risk and *CD38* expression levels and observed that high *CD38* expression conferred longer PFS across all risk categories, although statistical significance was not reached in the high-risk group due to limited sample size (**Figure 4C**). Collectively, these findings indicate that *CD38* gene expression serves as a predictive biomarker of PFS in NDMM patients treated with daratumumab-containing quadruplet regimens.

**Figure 4.**
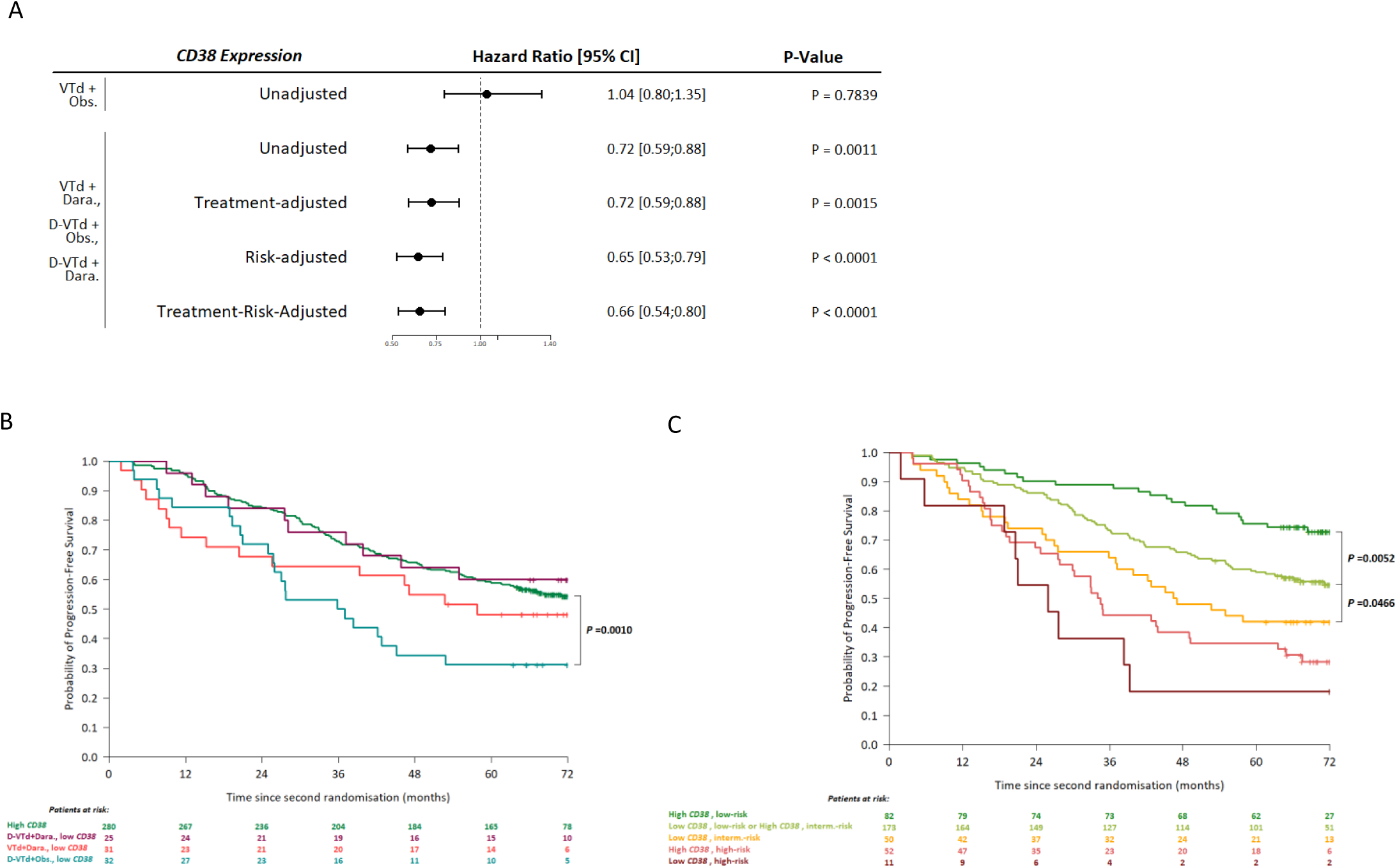
*CD38* expression analyses of PFS by RNA subtypes and risk status. (A) Forest plot displaying unadjusted and treatment- and/or risk-adjusted Cox models results (Hazard ratios, along with their 95% confidence intervals, and *P*-values) for patients according to daratumumab administration. (B) Kaplan- Meier survival curves for patients with daratumumab administration according to *CD38* expression and treatment arm: given that no significant difference was observed among the 3 daratumumab arms for patients with high *CD38* expression those patients were pooled (High *CD38*); logrank p-value for comparison of patients with high *CD38* expression vs. patients of D-VTd+Obs. arm with low *CD38* expression is displayed. (C) Kaplan-Meier survival curves for patients with daratumumab administration according to *CD38* expression and risk groups: given that no significant difference was observed between low *CD38/* Interm-risk patients and high *CD38*/Interm-risk patients those patients were hence pooled; logrank p-values for comparisons between high *CD38*/low-risk patients and low *CD38*/low-risk patients on one hand, and high *CD38*/Interm-risk patients and low *CD38*/Interm-risk patients on the other hand, are indicated.

### MRD-negativity rates according to transcriptomic risk status

D-VTd significantly improved MRD negativity rates post–autologous stem cell transplantation (ASCT) compared with VTd, irrespective of transcriptomic risk group. Unexpectedly, patients in the high-risk group (MS+MF subtypes) who received D-VTd achieved the highest MRD negativity rate (85.1%), exceeding that of the intermediate-risk group (CD+HP subtypes; 62.1%) (**Figure 5A**). This observation suggests that high-risk patients may initially reach deeper responses but experience earlier relapse compared with other subgroups, in contrast to the more sustained responses seen in intermediate-risk patients. Importantly, MRD negativity rates remained significantly higher in the high-risk group than in the intermediate-risk group among patients receiving daratumumab-containing regimens (*P* = 0.0154 after six months of maintenance), implying that relapse in this population may occur later rather than immediately (**Figure 5B**). In line with previous findings from CASSIOPEIA showing that MRD negativity at day 100 post-ASCT correlates with improved PFS,^17^ we examined this association across transcriptomic risk categories in patients treated with daratumumab during either phase 1 or phase 2 of therapy. A strong correlation between post-consolidation MRD negativity and superior PFS was observed in intermediate-risk groups (*P* < 0.0001) as well as superior OS (*P*=0.0040) (**Figure 5C–D**). Among low-risk patients, MRD-negativity was significantly associated with PFS (*P*=0.0023) but not with OS. As anticipated, MRD negativity was not associated with PFS or OS in the high-risk group. Taken together, these findings indicate that MRD status alone may not serve as a universally reliable predictor of survival across all molecular subtypes. This underscores the importance of integrating baseline omic data, particularly transcriptomic information, when interpreting MRD dynamics in MM.

**Figure 5.**
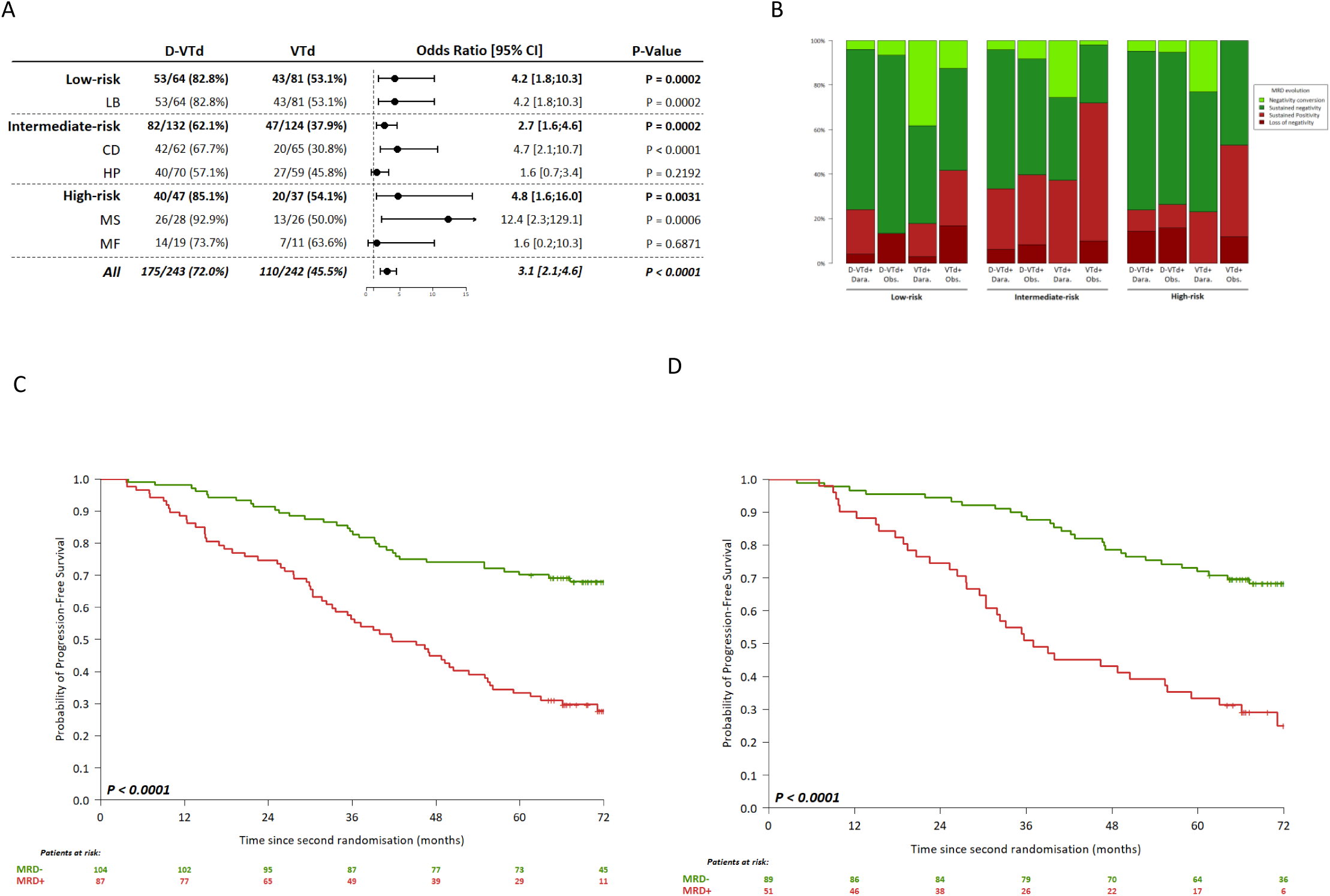
MRD-negativity rates according to transcriptomic risk status. (A) MRD-negativity rates at day 100 after ASCT according to RNA subtypes and risk status. Forest plot displaying comparison results for MRD according to first randomisation treatment arms in the different RNA subtypes and associated risk-groups. (B) MRD status evolution between day 100 after ASCT and 6 months post-consolidation according to risk status and treatment arm. Kaplan-Meier PFS curves for intermediate-risk patients with daratumumab administration, (C) according to day 100 after ASCT MRD satus and, (D) according to 6 months post-consolidation MRD satus.

## Discussion

Although the CASSIOPEIA trial demonstrated that patient survival was markedly improved with daratumumab-containing regimens, clinical outcomes remain heterogeneous. There remains an unmet need for patients at high risk of early disease progression. In this study, we assessed whether baseline tumor transcriptomic profiles could enhance risk stratification and provide biological context for the interpretation of MRD status evaluated by MFC, whose prognostic relevance for PFS and OS has been established in the overall cohort.

The results presented here represent the first risk-subgroup analysis based on transcriptomic profiling of NDMM patients treated with daratumumab, bortezomib, thalidomide, and dexamethasone. Using a large RNA-seq dataset from 628 CASSIOPEIA patients, we defined five transcriptomic subtypes and identified three risk groups associated with distinct long-term outcomes: 72-month PFS rates of 59.7%, 42.1%, and 23.3%, and corresponding OS rates of 83.3%, 87.8%, and 66.1% for the low-, intermediate-, and high-risk groups, respectively. Building on our previous findings that established the quadruplet therapy as the standard of care for transplant-eligible NDMM patients,^6–8^ we evaluated transcriptomic-based risk prediction among individuals who had received daratumumab at any treatment phase. Risk stratification derived from gene-expression clustering was preserved in these patients and consistently demonstrated the benefit of daratumumab across all risk categories. Furthermore, high *CD38* expression level was associated with better outcome of patients who received daratumumab regardless of treatment arms and transcriptomic risk.

Integrated analysis of MRD across the three risk groups **r**evealed strikingly divergent therapeutic trajectories associated with distinct transcriptomic signatures. Despite achieving deep responses and high post-consolidation MRD negativity rates, high-risk patients displayed markedly inferior PFS (26.6% at 72 months). In this group, gene-expression profiles were enriched for activation of the MAF transcription factor family (*c*-MAF, *MAFA*, *MAFB*), defining the MF subtype (5.9%). Importantly, the transcriptomic approach captures all patients with MAF family activation, extending beyond FISH-based cytogenetic classification (t(14;16), t(14;20)), which identifies only ∼80% of such cases. MF subtype is known to harbor chromothripsis and APOBEC mutational signatures, genomic hallmarks associated with poor prognosis^18–20^ and genomic instability.^21^ We can assume that although these cells are sensitive to induction/consolidation therapy, their capacity for rapid adaptation linked to genomic instability, potentially coupled with a hostile hypoxic and immunosuppressive remodeled microenvironment, favors the emergence of resistant clones, explaining the rapid relapse observed. This highlights the importance of incorporating microenvironmental analyses to better elucidate tumor–immune environment interactions. However, regarding MF subtype, MRD results must be interpreted with caution, firstly for reasons of statistical power, as the subtype sample size is small, and secondly due to potential limitations in the measurement of MRD by MFC.^22^ The most significant limitation is the known association between extramedullary disease and MF subtype.^23^ For these patients, in particular, it appears necessary to measure the effectiveness of treatment in the entire bone medulla and extramedullary compartments, FDG PET/CT response can provide complementary information to assessment of MRD by MFC.^24^

The high-risk group also included patients with *NSD2* overexpression resulting from the t(4;14) translocation. *NSD2*-driven expansion and redistribution of H3K36me2 reshapes 3D chromatin architecture,^25^ leading to the activation of oncogenic pathways that may underlie poor outcomes.^26^ Despite significant therapeutic advances, high-risk patients continue to relapse and should be prioritized for clinical trials investigating agents with novel mechanisms of action, such as therapies targeting metabolic vulnerabilities driven by oncogenic transcriptional programs^27^ or T-cell–redirecting bispecific antibodies.^28^ Interestingly, expression levels of immunotherapy targets vary among subtypes: all four major targets are highly expressed in the MS subtype, whereas *GPRC5D* is most strongly expressed in the MF subtype, potentially representing an optimal bispecific T-cell engager target in this group (**Supplemental Figure 10**). In contrast, the low-risk group, characterized by a high dormancy gene signature,^15^ exhibited exceptional post-consolidation MRD negativity (82.8%) and durable PFS (70.2% at 72 months). Elevated dormancy scores correlated with improved outcomes, consistent with previous findings.^15^ Overexpression of *AXL* suggests increased dependence on the bone marrow microenvironment, possibly through the AXL–GAS6 axis, which maintains myeloma cell dormancy within the endosteal niche.^15,29–30^ Given that low-risk patients are more likely than others to have drug-resistant dormant cells that may be reactivated by microenvironmental cues, they could benefit from lighter maintenance regimens with early discontinuation in the setting of sustained MRD negativity, potentially combined with agents that prevent exit from dormancy. Between these extremes, the intermediate-risk group, encompassing more than half of patients, those overexpressing *CCND1* or exhibiting hyperdiploidy, was characterized by moderate MRD negativity rates (62%) and slower progression, suggesting a balance between proliferation and disease control. In this group, MRD status was strongly correlated with both PFS and OS, supporting dynamic MRD reassessment to better capture MRD kinetics and guide maintenance therapy optimization.^31^

As induction and consolidation therapies evolve, these results highlight the importance of risk-adapted treatment and longitudinal MRD evaluation.

## Acknowledgments

This study was sponsored by the Intergroupe Francophone du Myélome and supported by the Dutch- Belgian Cooperative Trial Group for Hematology Oncology, Johnson & Johnson, the Région Pays de la Loire (Connect Talent program, RPH2114NNA), Nantes Métropole (RPH2114NNA), CHU Nantes, the I-SITE NexT (ANR-16-IDEX-0007), HéMA-NExT research cluster and the SIRIC ILIAD (INCa-DGOS-INSERM-ITMO Cancer_18011). We thank the laboratory staff who processed the samples at the UGM, IUC- T Oncopole, Toulouse, and the Genomics and Bioinformatics core facility of Nantes (GenoA, BIRD, Biogenouest, IFB) for its technical support. We warmly thank all of the CASSIOPEIA study participants and their families for making this research possible.

## Authorship Contributions

S.M. and P.M. conceived the study; F.M. C.G.-C., and S.M. analysed and interpreted the data; V. B.-G, A.P., C.T., M.v.D., Magali D., E.B., E.L., P.S., J.C. acquired or collected the data; F.M. and S.M. wrote the manuscript. C.G.-C., E.L., Marie D., N.G., P.M. contributed to the paper writing and revision. All authors have read and agreed to the published version of the manuscript. All authors had full access to all the data in the study and had final responsibility for the decision to submit for publication.

## Data sharing statement

Data are available on request from the corresponding authors Stéphane Minvielle and Philippe Moreau.

