## Supplemental information for "Expression profile of CASSIOPEIA patients refines prognostic value of MRD negativity in multiple myeloma"

### Expression profile of CASSIOPEIA patients refines prognostic value of MRD negativity in the area of quadruplet therapy in multiple myeloma

Magrangeas *et al.*

**Supplemental Table 1:** Characteristics of patients at baseline

**Supplemental Figure 1:** Trial profile of the RNA-seq cohort

**Supplemental Figure 2:** Progression-free and Overall Survivals according to treatment arm in the RNAseq cohort

**Supplemental Figure 3:** RNA-seq consensus clustering matrix

**Supplemental Figure 4:** Boxplot displaying distribution of the score for PR group according to the 5 RNA subtypes

**Supplemental Figure 5:** Gene ontology pathway enrichment investigated for MF, MS, HP, CD and LB RNA subtypes one vs. others

**Supplemental Figure 6:** Unsupervised hierarchical clustering of the 40-gene dormancy signature of the CASSIOPEIA RNAseq cohort

**Supplemental Figure 7:** RNA subtypes characteristics

**Supplemental Figure 8:** *CD38* expression comparison between the fourth treatment groups

**Supplemental Figure 9:** PFS according by *CD38* expression in patients who received daratumumab-containing quadruplet regimen

**Supplemental Figure 10:** Gene expression levels of immunotherapy targets in RNA subtypes of CASSIOPEIA patients

| Parameter | Transcriptomic cohort (n=589) | Other patients (n=496) | P-value |
|---|---|---|---|
| ***Age*** | | | *NS* |
| *Mean (sd)* | 56.8 (7.0) | 56.4 (7.0) | |
| *Median (Range)* | 59 (26;65) | 58 (22 ;65) | |
| *Missing* | *0* | *0* | |
| ***Sex*** | | | *NS* |
| Male | 339 (57.6%) | 296 (59.7%) | |
| Female | 250 (42.4%) | 200 (40.3%) | |
| *Missing* | *0 (-)* | *0 (-)* | |
| ***t(4 ;14)*** | | | *NS* |
| No | 519 (89.0%) | 381 (90.5%) | |
| Yes | 64 (11.0%) | 40 (9.5%) | |
| *Missing* | *6 (-)* | *75 (-)* | |
| ***del(17p)*** | | | *NS* |
| No | 536 (91.9%) | 387 (91.9%) | |
| Yes | 47 (8.1%) | 34 (8.1%) | |
| *Missing* | *6 (-)* | *75 (-)* | |
| ***Risk*** | | | *NS* |
| High risk | 97 (16.5%) | 71 (14.3%) | |
| No high-risk factor identified | 492 (83.5%) | 425 (85.7%) | |
| *Missing* | *0 (-)* | *0 (-)* | |
| ***ISS*** | | | *0.0006* |
| Stage I | 212 (36.0%) | 235 (47.4 %) | |
| Stage II | 278 (47.2%) | 194 (39.1%) | |
| Stage III | 99 (16.8%) | 67 (13.5%) | |
| *Missing* | *0 (-)* | *0 (-)* | |

**Supplemental Table 1. Characteristics of patients at baseline.** Demographics and cytogenetic parameters according to RNA-seq data availability, for patients who went through at least the first randomisation.

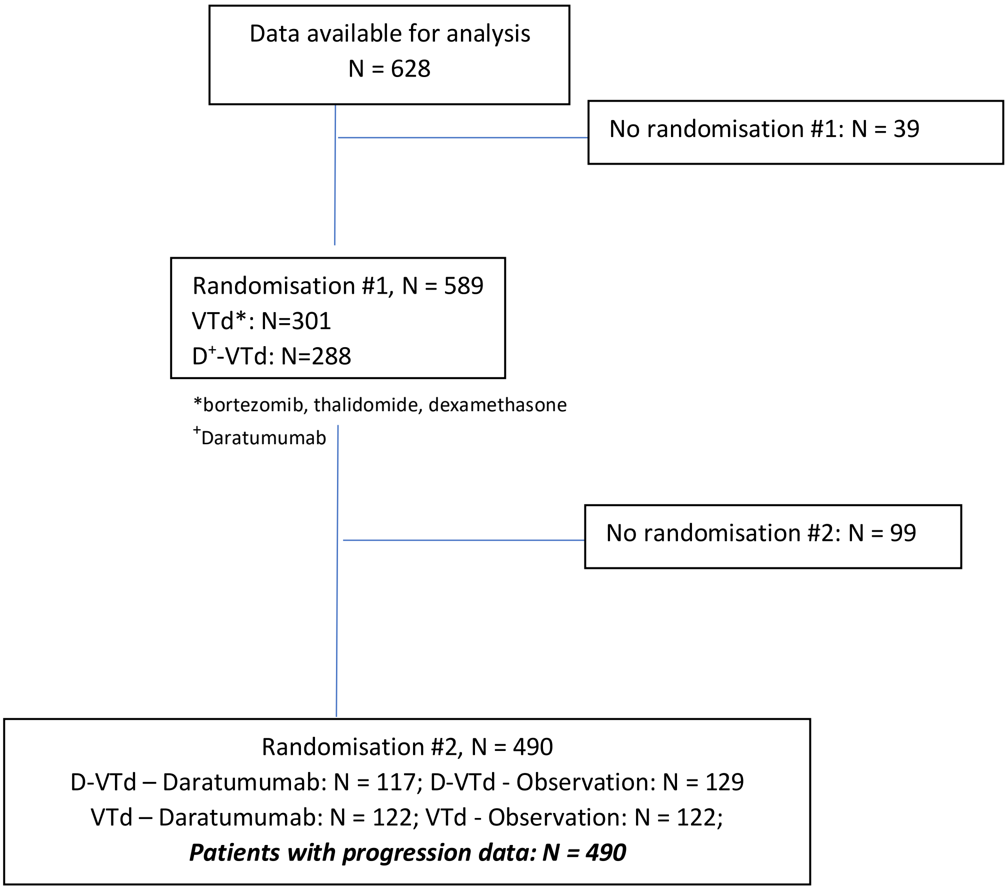


**Supplemental Figure 1**. **Trial profile of the RNA-seq cohort.** Flowchart representing patient distribution at the different timepoints of the study, for patients with available RNA-seq data.

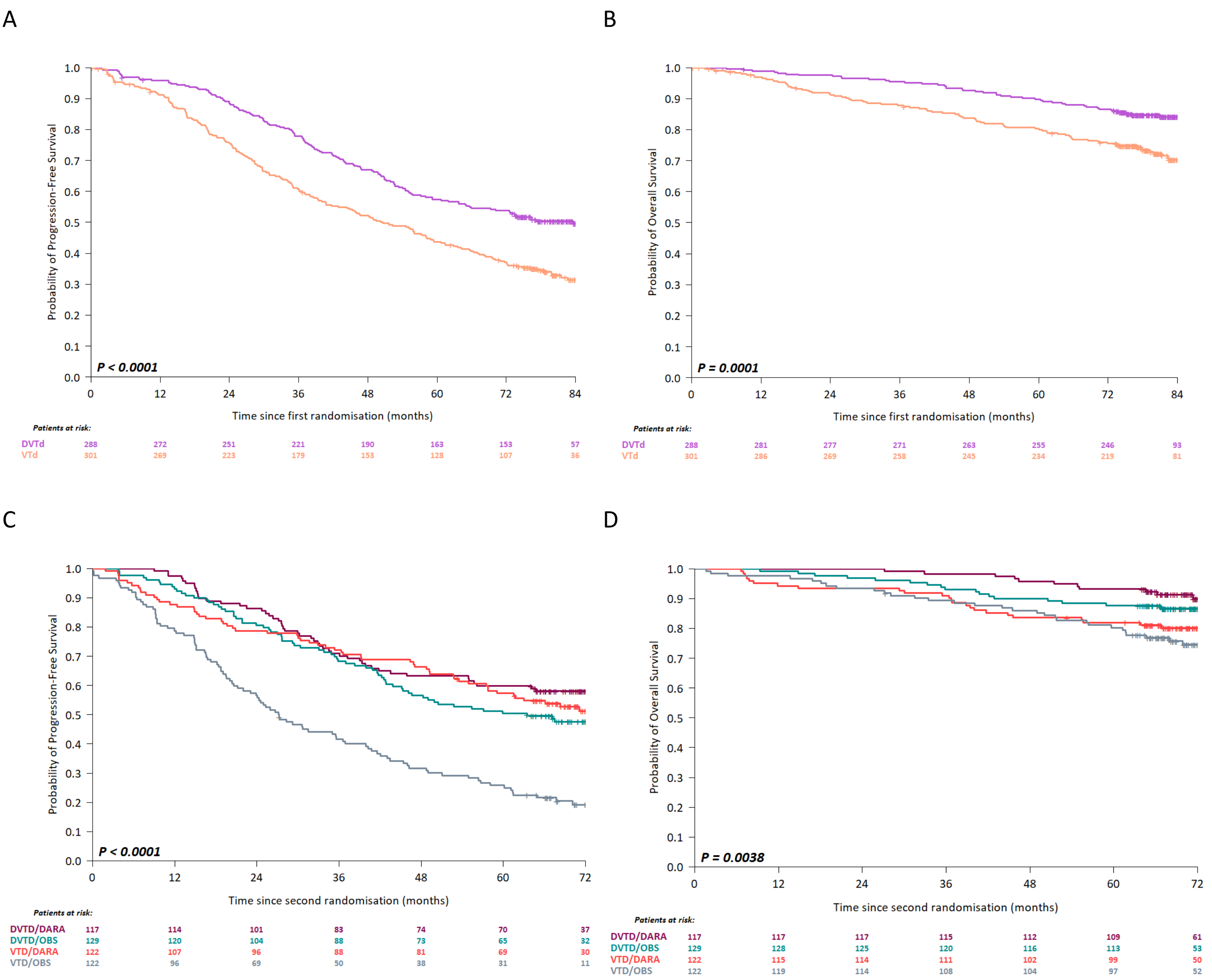


**Supplemental Figure 2. Progression-free and Overall Survivals according to treatment arm in the RNAseq cohort.** (A) Progression-Free Survival from first randomisation according to treatment. (B) Overall Survival from first randomisation according to treatment. (C) Progression-Free Survival from second randomisation according to treatment arm; pairwise comparisons identified two significantly different subgroups D-VTd + Dara., D-VTd + Obs., VTd + Dara. and VTd + Obs. (*P* < 0.0001). (D) Overall Survival from second randomisation according to treatment arm.

A B

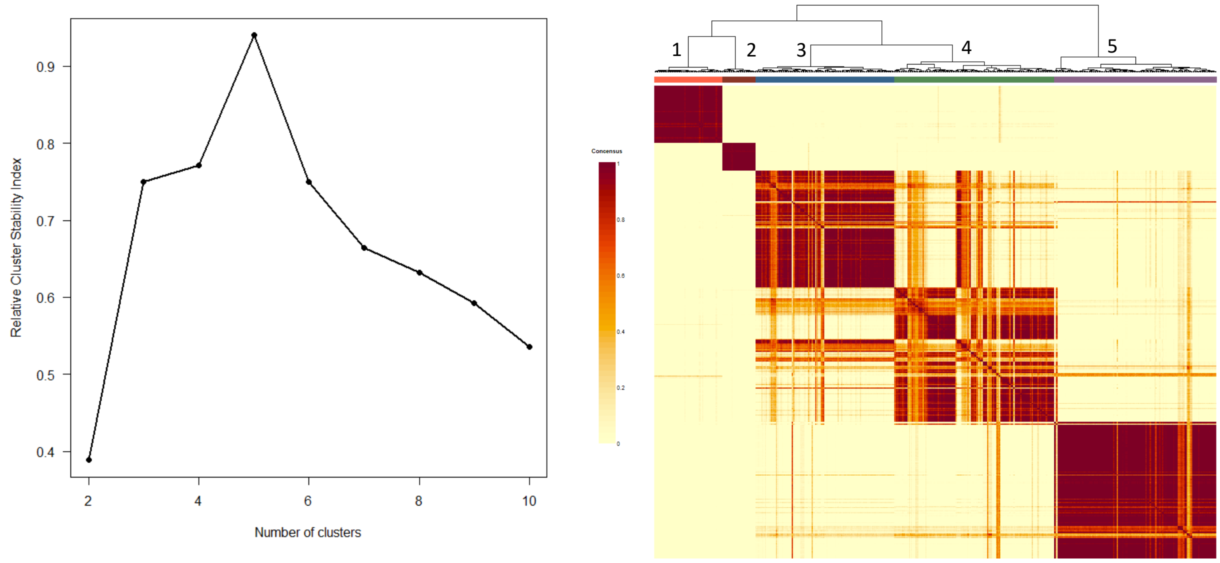


**Supplemental Figure 3**. **RNA-seq consensus clustering matrix**. (A) Optimal number of clusters determination from gene expression consensus clustering based on relative cluster stability index. (B) RNA-seq consensus clustering matrix. Consensus matrix showing the consistency of class assignment for K = 5 clustering of RNA-seq data derived from 628 samples and 932 feature-selected genes.

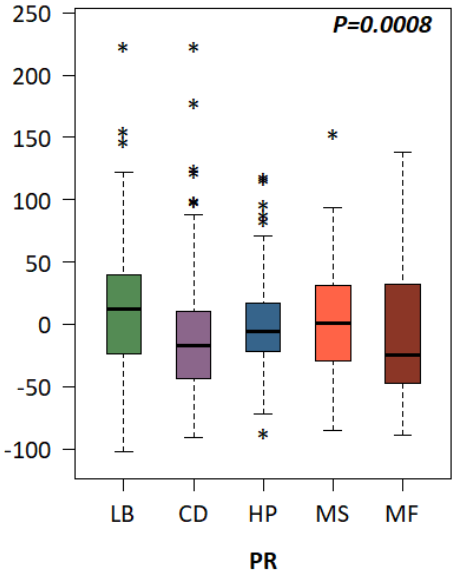


**Supplemental Figure 4. Boxplot displaying distribution of the score for PR group[10] according to the 5 RNA subtypes.**

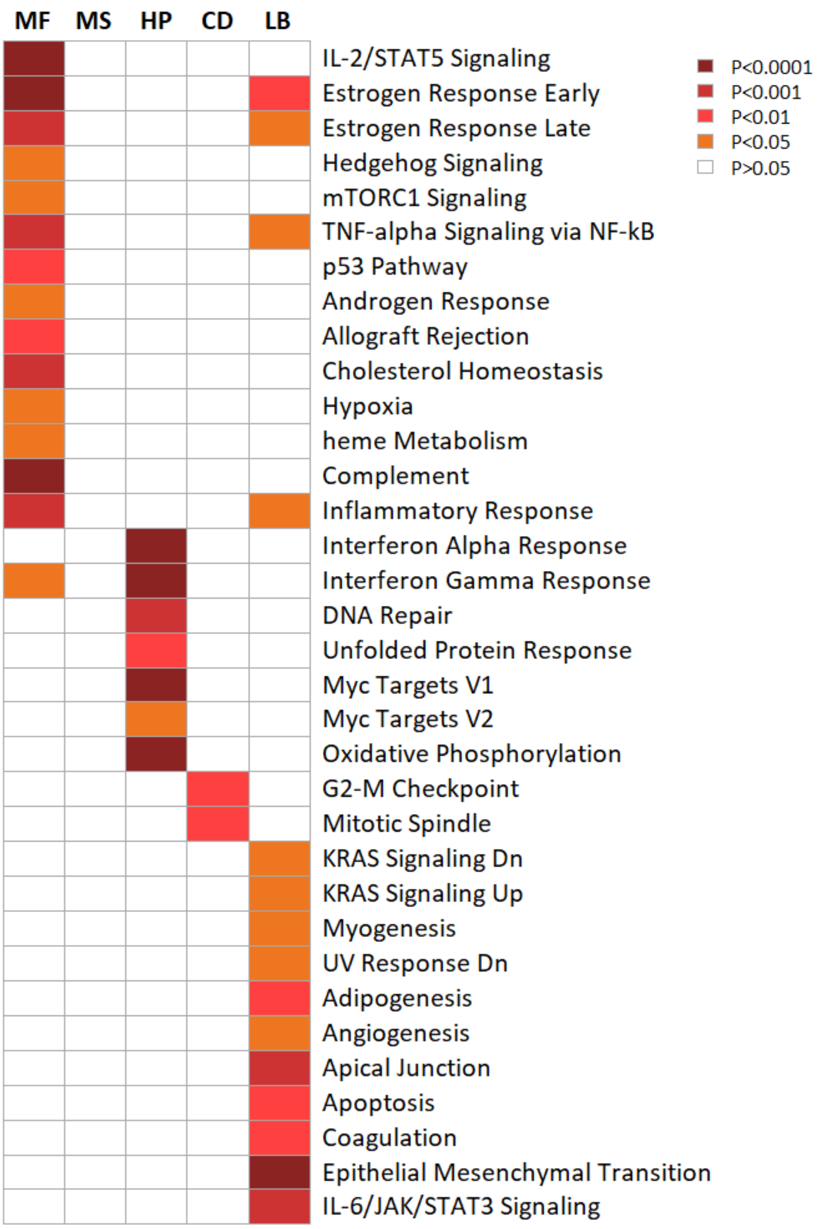


**Supplemental Figure 5. Gene ontology pathway enrichment investigated for MF, MS, HP, CD and LB RNA subtypes one vs. others.** Hallmark gene sets from MSigDB Human collections was used to test pathway enrichment

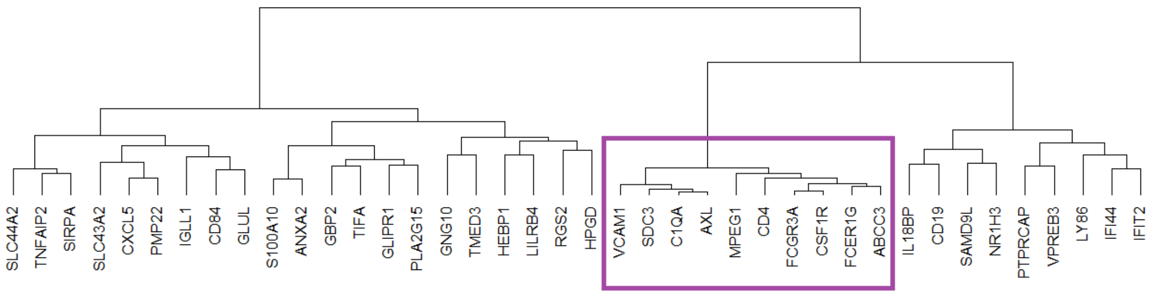


**Supplemental Figure 6. Unsupervised hierarchical clustering of the 40-gene dormancy signature[15] of the CASSIOPEIA RNAseq cohort.** The gene dendrogram identified *AXL*-coregulated signature, the branch containing *AXL*-coregulated signature is boxed.

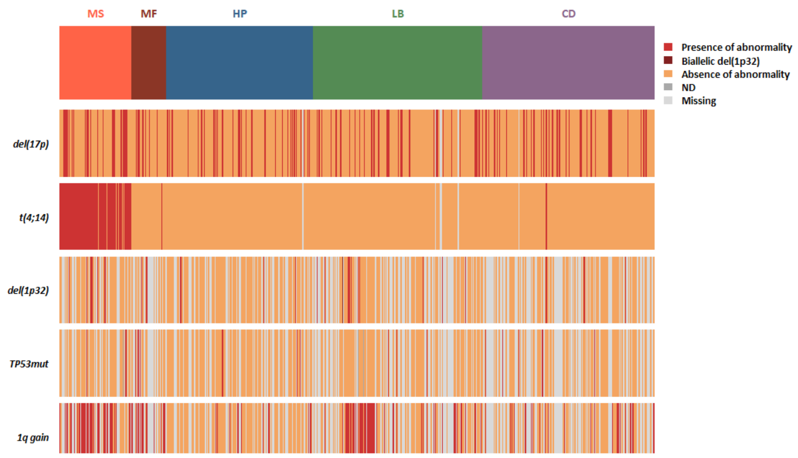


**Supplemental Figure 7. RNA subtypes characteristics.** RNA subtypes and genomic high-risk markers: del(17p) with a cutoff of > 20%, t(4;14), del(1p32), *TP53* mutation and 1q gain.

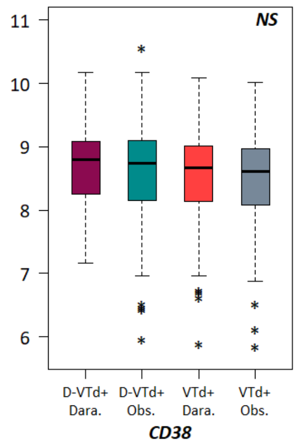


**Supplemental Figure 8. *CD38* expression comparison between the four treatment arms.** Boxplots displaying *CD38* expression distribution according to treatment arms.

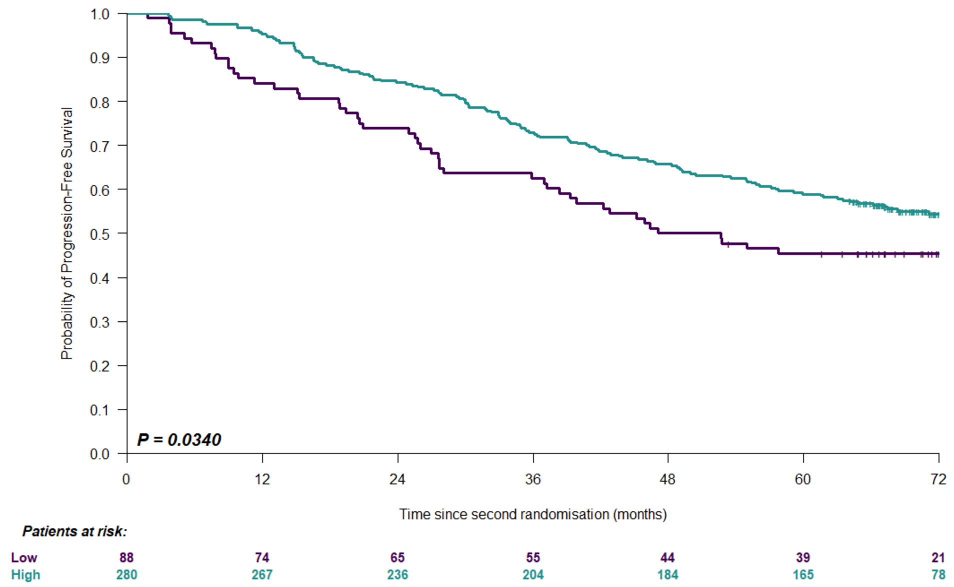


**Supplemental Figure 9. PFS according by *CD38* expression in patients who received daratumumab-containing quadruplet regimen**. Kaplan-Meier curve and logrank test result for PFS according to *CD38* expression, high *vs.* low. High expression of *CD38* was defined as quartiles 2 to 4 and low as quartile 1.

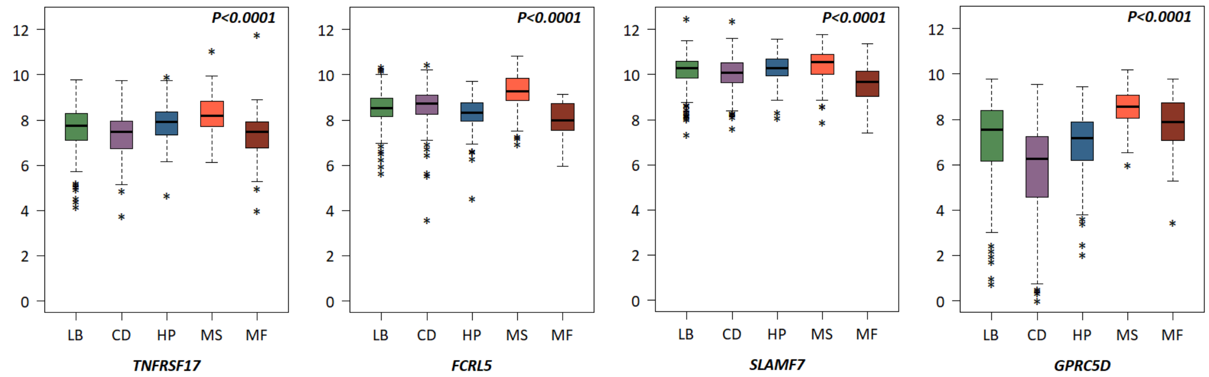


**Supplemental Figure 10. Gene expression levels of immunotherapy targets in RNA subtypes of CASSIOPEIA patients.** Boxplots representing expression levels of the 4 current T-cell immunotherapy targets *TNFRSF17*, *FCRL5, SLAMF7* and *GPRC5D* in the five RNA subtypes.
